# MillionFull enables low-cost, massive, full-length enzyme sequence–fitness data collection for machine learning–guided enzyme engineering

**DOI:** 10.1101/2025.10.24.684421

**Authors:** Jinbei Li, Bjarke Erichsen, Simon R. Krarup, Sonia C. Yuan, Kenan Jijakli, Søren Karst, Lei Yang, Alex Toftgaard Nielsen

## Abstract

Machine learning holds great promise for accelerating enzyme optimization, but its power is fundamentally constrained by the limited availability of sequence–fitness data. Here, we introduce **MillionFull**, a low-cost method that enables high-throughput full-length sequence– fitness mapping for enzymes of arbitrary length. Each run yields on the order of 10⁵–10⁷ data points, capturing sequence–function relationships at unprecedented scale. By overcoming the data bottleneck, MillionFull provides a foundation for dramatically advancing AI-driven enzyme engineering.

## Introduction

The use of machine learning holds great promise for accelerating enzyme optimization^1,2^, but this approach is fundamentally constrained by the availability of data^1–3^. Conventional data collection approaches for enzyme-activity, relying on individual enzyme variant measurements, is tedious and costly, even with the use of robotic platforms.

The adoption of ultra-high throughput screening approaches based on the use of droplet sorting or growth-coupled assays is growing. A great amount of enzyme activity information is embedded in such workflows. In droplet sorting, the information takes the form of variant distribution among the two or several sorted bins^4,5^; in growth-coupled assays, the information is in the form of the growth rate of the population of cells containing a particular enzyme variant. The latter has the advantage of higher resolution (a continuous readout within the dynamic range), and lower noise.

Extracting such rich information is not straightforward. While deep sequencing can capture the dynamics of the changing distribution of DNA sequence variants (the basis from which growth rate can be derived) over the course of a growth-based assay^6,7^, it is generally limited to 300 or 600 bp of read lengths, while most enzyme-encoding genes are close to or longer than 1 kb. While long-read sequencing can easily cover the full length of most enzyme-encoding DNA, it suffers from low read accuracy and throughput^8,9^.

We therefore developed MillionFull, a method that combines growth-based selection, deep sequencing, and long-read sequencing to generate full-length sequence-fitness datasets at unprecedented scale (**Fig. 1A, B**). The key to achieving this combination is unique molecular barcoding of the target enzyme-coding sequences during library creation. First, the full-length sequence-barcode region of the unselected library is sequenced with long-read sequencing. This is done for two purposes: 1) correcting sequencing errors through consensus generation of the reads with the same barcodes; 2) generating a full-length sequence-to-barcode lookup table. Second, the barcode region of samples taken during selection is sequenced through deep sequencing to profile growth rates of variants. The barcodes are then used to link growth rates, a proxy for target enzyme fitness, to the corresponding full-length sequences, resulting in rich sequence-fitness relationship data. The throughput of MillionFull is largely limited by the sequencing throughput and accuracy of long-read sequencing (constrained by the number of redundant reads required to accurately resolve a sequence). For example, with a read throughput of 10^8^ and redundancy requirement of 10-30 reads, up to ∼10^7^ data points can be collected.

**Fig. 1.**
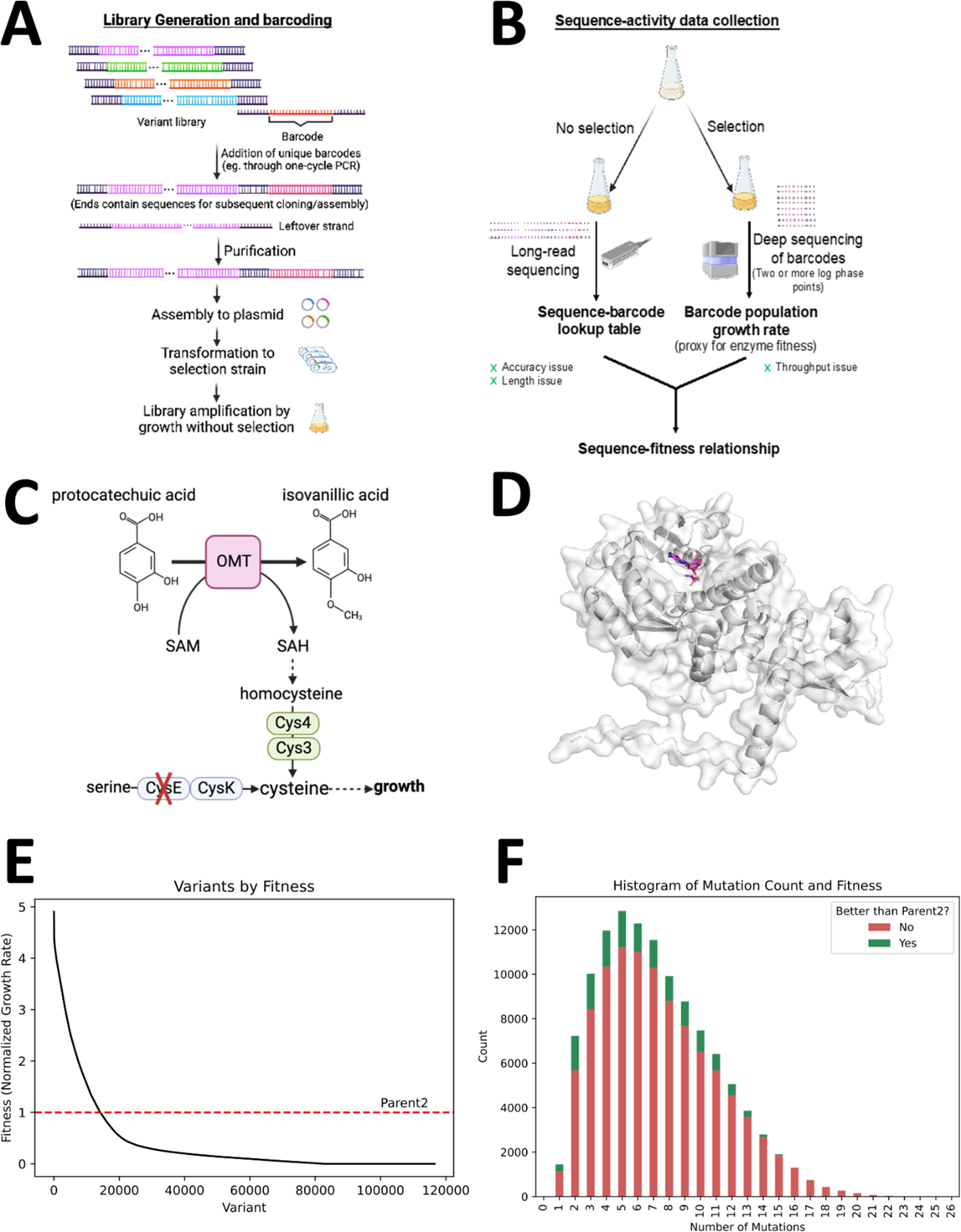
MillionFull workflow and scale. **A.** Library generation and barcoding **B.** Sequence-fitness data collection. **C**. selection system used for validation of the MillionFull method. **D.** AlphaFolded structure of AtOMT1. The cofactor SAH is shown in magenta. **E.** Normalized growth rate of unique protein variants, sorted by fitness. The red dashed line is the Parent2 fitness. **F.** Histogram of the number of amino acid substitutions. Variants with a fitness higher than Parent2 are colored green, variants with fitness lower than Parent2 are colored red.

## Results

### Growth-Coupled Selection System

The system we used in this study to demonstrate MillionFull was a previously established growth coupling system where *E. coli* cell growth is coupled to the activity of a S-adenosyl methionine (SAM)-dependent O-methyltransferase (MTase)^10^. In this system, the methylation byproduct, S-Adenosyl-L-homocysteine (SAH), serves as a precursor for cysteine while the endogenous cysteine synthesis pathway is disrupted (**Fig. 1C**). This design allows the use of growth rate as a proxy for MTase activity under selective growth conditions. We used protocatechuic acid (PCA) as the substrate to be methylated by the target MTase (depending on the methylation site, the product is either vanillic acid or isovanillic acid, **Fig. 1C**).

### Library Generation

AtOMT1 (EC:2.1.1.42, GenBank: AAB96879.1)^1^, a methyltransferase that showed activity for producing only isovanillic acid from PCA, was chosen as the initial template for variant library generation (**Fig. 1D**). Because the initial activity was low, to increase both data diversity and the amount of data points with significant activity, we first created an error-prone PCR (epPCR) library from wild-type AtOMT1 (Parent1), subjected it to selection with the MTase growth coupling system, and used the resulting selected variant mixture to serve as the template for eventual epPCR library creation and data extraction.

### Barcoding and Library Amplification

A random DNA barcode fragment of 25 bp (>10^15^ possibilities) was attached to the coding sequences using a one-cycle PCR and gel purification procedure. The barcoded DNA library was assembled onto an *E. coli* expression vector and transformed to the selection strain. A portion of the library, estimated based on colony counting to consist of 1.5 million variants (this turned out to be an underestimate given subsequent deep sequencing results), was taken and amplified by growth without selection, serving as the *in vivo* library for the following steps.

### Nanopore Sequencing and Deep Sequencing

Nanopore sequencing was performed on the *in vivo* library after non-selective growth amplification, generating 19.3 M reads. The distribution of variant abundance during growth-based selection was captured using deep sequencing of samples taken at 6 time points including one taken before the selection started, with 20-33 M reads for each sample. As expected, the distribution of read counts for different barcodes became increasingly uneven as selection progressed (**Supplementary Fig. 1**). We used this distribution in combination with OD measurements to calculate a growth rate for each variant using the DiMSum pipeline^11^.

Deep sequencing revealed the actual library size to be ∼3.2 M variants as judged by the number of unique barcodes with more than 5 reads in total at time point 0. Rather than expending extensive Nanopore sequencing efforts on the unselected library to capture all sequence-barcode relationships, we performed an additional Nanopore sequencing run (1.5 M reads) on the post-selection library. This biases the data collection toward higher-fitness variants, which are more informative^12^. After the additional Nanopore data is added, all but one of the top 1000 barcodes obtained at least 7 reads - the threshold used for full-length consensus sequence generation.

### Data Size and Features

Since the template used to generate the second library for data collection was a mixture, we defined the most common sequence in the unselected second library as “Parent2”, containing six point mutations compared with Parent1 (see **Extended Data 8** for sequences). By merging calculated growth rates for Parent2-identical variants, all variants were also assigned a fitness value relative to the Parent2 enzyme.

Over 571,000 data points of unique barcodes correlating full-length DNA sequences to fitness were collected (based on aggregate fitness values from all time points) (**Supplementary Fig. 2**). Indels and premature stop codons in the coding sequence were common, likely due to the high-error rate conditions used during epPCR, and the fact that the sequences come from two rounds of epPCRs. More than 100,000 data points, termed “meaningful data”, remained after excluding sequences with indels, premature stop codons before the last active site (residue index 327, with full length being 363), or mutations in the promoter/RBS region.

Among the MillionFull data points collected, over 12,000 unique DNA sequences have a predicted fitness higher than the Parent2 sequence (p<0.01, based on 4 different selection time points). Interestingly, the data revealed a group of around 30 sequences which have a section of 61 AAs deleted from AA102-162 yet, generally, showed improved fitness. This finding further highlights the importance of capturing the full length of the enzyme sequence variants, as such deletions may be more easily missed by deep sequencing-only approaches.

Correlations in sequence-fitness mappings among the triplicates were all above 0.91, indicating a low noise level in the data (See DiMSum report in **Extended data 1**). Fitness levels for variants with premature stop codons mostly center around 0, and fitness levels for variants with Parent2-identical sequence are generally within 0.5-1.5 (**Supplementary Fig. 3**).

The sequences of the meaningful data are highly diverse (**Supplementary Fig. 4**). There are ∼116,000 unique protein sequences without indels or early stop codons, and the library has an average of 5 amino acid substitutions per sequence, ranging up to a maximum of 26 substitutions. Even among variants with a fitness score above the Parent2-level, most sequences have 2-10 and up to 18 AA-level mutations (**Fig. 1D**). Additionally, there is good coverage of amino acid mutations across all 363 positions of the protein; most residue positions have sampled over 10 of the 19 possible mutations (**Supplementary Fig. 4C**).

### Data Accuracy

We estimated the error rate of consensus sequences with a given number of reads by subsampling that number of reads from Nanopore read clusters with large numbers of reads. Seven reads, the chosen threshold for the final data, gave an error rate of 3.56% for the entire consensus coding sequence compared with the consensus sequences based on the entire clusters, or 0.0033% per bp. Note that most of the variants in the final data had more than seven Nanopore reads. 41 sequences, including 9 with indels, were amplified from the library using their barcodes as one of the primers, and sequenced by Sanger sequencing. In all 41 cases, Sanger sequencing results matched 100% with the consensus sequences based on Nanopore reads (see **Extended data 2** for their cluster size information).

The accuracy of the MillionFull sequence-fitness mappings was empirically validated through individual measurements made on a set of ∼30 variants spanning the full fitness range. Individual growth rates measured in the selection condition, and enzyme activities measured through lysate-based bioconversion of PCA to IVA both show strong positive correlation (**Fig. 2A, B**).

**Fig. 2.**
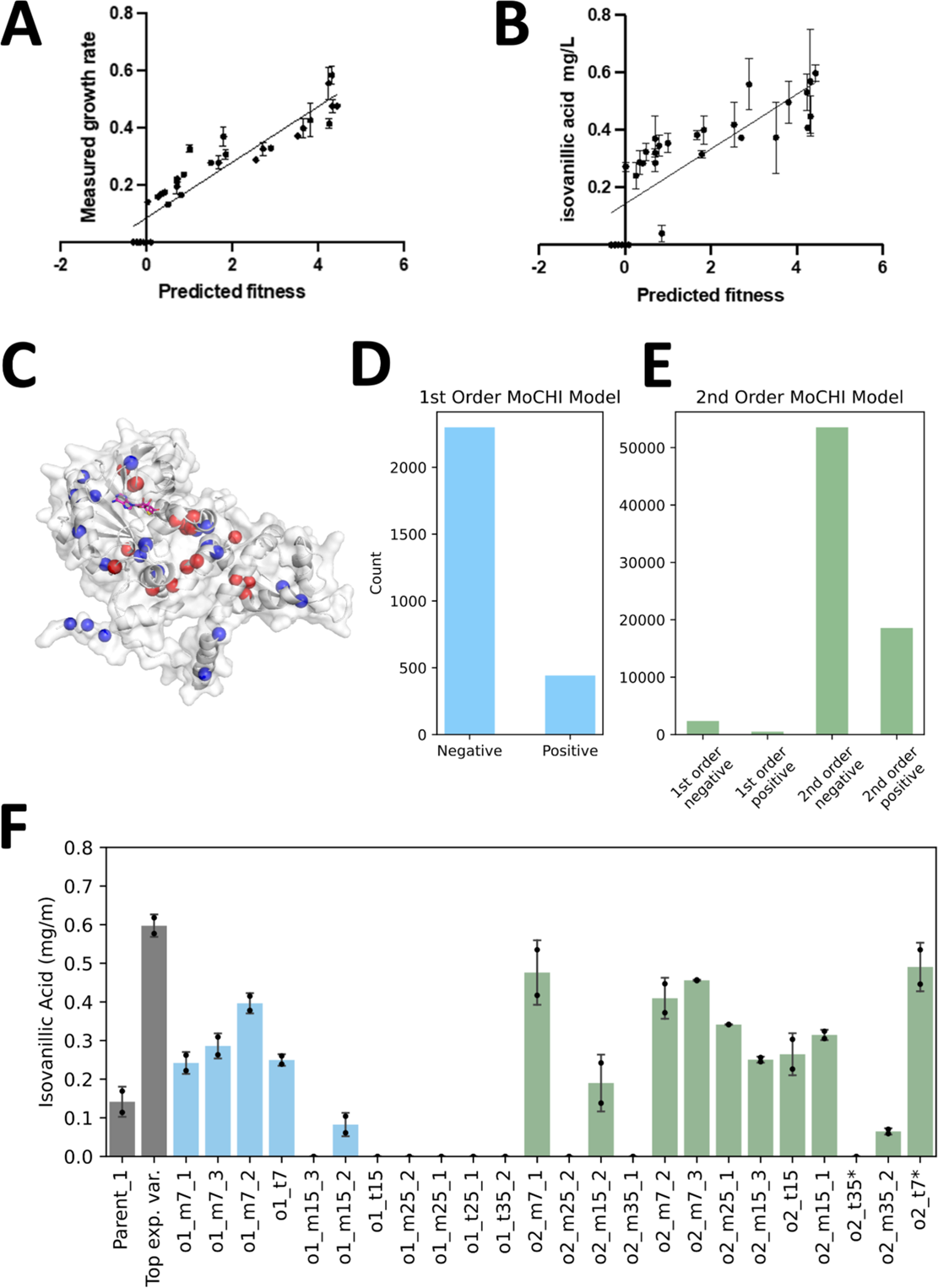
Empirical data validation and learning. **A.** Correlation between individually measured growth rates and fitness levels in the MillionFull data (R^2^ = 0.87). **B.** Correlation between individually measured enzyme activities in bioconversion assays and fitness levels in the MillionFull data (R^2^ = 0.65). **C.** The top 20 positions with the most deleterious (red) or beneficial (blue) point mutations. SAH is shown in magenta. **D.** Positive and negative point mutations identified by 1st order model. **E.** Positive and negative point mutations and epistatic interactions identified by 2nd order model. **F.** Isovanillic acid production of MoCHI-designed variants. Controls are in grey, variants designed via the first-order model and filtered with ESM & ProteinMPNN are in blue, and variants designed via the second-order model are in green. “o1” and “o2” indicate the MoCHI model; “mx” indicates the variant was designed via stochastic mutation sampling with x mutations, and “tx” indicates the variant was designed by deterministically combining the top mutations or mutation pairs with x mutations; the last number is the code for variants in the same category. o2_t7 and o2_t35 had a G2D mutation and a D2G mutation, respectively. They are labeled with an asterisk (*) to indicate the deviation from the original design.

### Learning the Effects of Point Mutations and Epistatic Interactions

We used MoCHI^13^, a tool for using neural networks to fit interpretable models for quantifying the effects of mutations and their epistatic interactions (**Supplementary Fig. 5**). The resulting 1^st^ order model identified 442 mutations with positive effects, and 2300 mutations with negative effects (**Fig. 2D**). We mapped the positions where mutations with the most aggregate positive or negative effects were observed (top positive/negative positions) (see **Extended Data 7** for mutation and position identities). While the top 20 negative positions are clustered around the active site, as expected, the top 20 positive positions are spread out across the enzyme’s structure with no obvious pattern. (**Fig. 2C**). This highlights the difficulty of rational selection of sites for mutagenesis during enzyme optimization. A 2^nd^ order model fitted using MoCHI additionally identified 18586 epistatic interactions with positive effects, and 53557 epistatic interactions with negative effects (**Fig. 2D**).

### Machine Learning-Guided Sequence Design

We leveraged the mutational effects learned by MoCHI to design novel sequences that were at least five mutations removed from all sequences in the data. This was done in two ways: 1) top point mutations were combined, either deterministically, or stochastically with higher likelihood for those with higher positive effects, and the resulting sequences were filtered through a combination of ProteinMPNN and ESM1v to minimize disruptive mutation combinations; 2) top point mutations were combined deterministically or stochastically, with a recursive sampling and evaluation loop to ensure compatibility among the chosen mutations based on their 2^nd^ order interaction effects. Despite the novelty of the sequences compared with any sequence in the data, multiple sequences, especially those designed based on the 2^nd^ order model, showed ∼3-fold increases in enzyme activity compared with Parent1, similar to the top variant from the screening data (**Fig. 2F**). Despite the additional pretrained model-based filtering, only 1 out of the 4 stochastically designed sequences with 15 or 25 mutations based on the 1^st^ order model showed detectable enzyme activity, being no higher than Parent1. In contrast, 4 out of 5 of the stochastically designed sequences with 15 or 25 mutations based on the 2^nd^ order model showed detectable enzyme activity, all showing a fitness score similar or higher than the Parent1 sequence. This further affirms the value of capturing epistatic interactions during enzyme fitness data collection.

## Discussion

Machine learning is a way to extract information, or patterns, from empirical data. While there has been an explosion of various learning approaches to extract information, the power of these efforts for protein learning is severely constrained by data availability. Two studies have examined training data size requirements for training deep learning models to predict protein-ligand binding, one using experimental datasets and the other physics simulation-based datasets. In both cases, model performance gains started to diminish only after a threshold of ∼10^5^ training samples was reached. This amount is several orders of magnitude larger than most existing protein sequence-fitness datasets^14,15^. Considering enzyme catalysis is more complex and dynamic than static binding interactions, it is reasonable to expect that the data requirement for effective enzyme fitness prediction using machine learning would be no less high.

The MillionFull approach offers a powerful and cost-efficient way to solve the data scarcity challenge in enzyme machine learning. With a labeled data output at a scale of 10^5^ – 10^7^, which will likely further expand with sequencing technology advancements, it has the potential to greatly expand the mapping of enzyme fitness landscapes, and boost the efficiency of machine learning-guided directed evolution.

During the preparation of this manuscript, several papers used or proposed similar workflows. Prywes et al. in 2025 reported the use of combining long-read sequencing, deep sequencing, and growth-based assay to profile the fitness landscape of rubisco^16^. Unfortunately, the data was limited to single point mutations, which can’t provide significant information on epistatic interaction effects. As rubisco is known to be highly constrained evolutionarily in the specificity vs catalytic rate trade-off^17^, and that single point mutations are much easier to access than multiple simultaneous mutations, it is unlikely that stepwise single point mutations could break the rubisco performance ceiling, while multiple simultaneous mutations as done in this work could provide a better chance.

The data collected in the present work is aggregate fitness effects, incorporated in which are enzyme kinetics, solubility, expression levels, stability. The Align Foundation recently proposed an approach in the GROQ-seq platform^18^ aimed at dissecting enzyme fitness parameters into k_cat_, K_m_, and enzyme abundance^19^. Compared with the MillionFull workflow proposed in the present work, GROQ-seq adds fitness measurements in varying substrate concentrations to obtain target enzyme K_m_ and v_max_, and separate fitness measurements of target protein-antibiotic resistance marker fusion proteins to obtain variant concentrations. Such additional assays would mean data output for less enzyme variants compared with a single assay under a given budget constraint, but provides deeper information for more mechanistic machine learning approaches and for greater adaptability of the learning outcome for diverse substrate concentrations and expression settings. In contrast, a single MillionFull runs afford project agility where the application settings are clearly defined.

Results from ongoing collaborations leveraging the reported dataset for advanced machine learning-based sequence design will be reported in future updates of this manuscript.

## Supporting information

Extended data-MillionFull

## Acknowledgments

We would like to express our sincere thanks to: Dr. Le Yuan, Dr. Feiran Li, Dr. Bruce Wittmann, Jason Yang, and Jonathan Funk for the valuable discussions that motivated the study; Prof. Paul Jensen and Dr. Viji Kandasamy for their contributions to the method design; Dr. André Faure and Dr. Arsenios Vlassis for their assistance with Illumina sequencing and data analysis; Dr. Linda Ahonen and Adrian Frey for their support with chemical analytics; Dr. Søren Karst, Dr. Scott Quinoo, Troels Hansen, Dr. Tue Jørgensen, Dr. Zofia Jarczyńska, and Keyan Liu for their contributions to nanopore experiments and data analysis; Dr. Se Hyeuk Kim, Dr. Christoffer Rode, Christina Lenhard, and Lena Heer for their help with molecular biology experiments. This work was funded by the Novo Nordisk Foundation grant NNF20CC0035580 and Enzidia ApS.

## Data Availability

Growth rates for each DNA sequence and normalized fitness values per protein sequence are uploaded at https://doi.org/10.5281/zenodo.17282389.

## Materials and Methods

### Media, growth and transformation conditions

During strain construction and maintenance, *E. coli* strains were grown at 37 °C in LB or 2x YT, supplemented with 50 mg/L L-cysteine for the selection background strain, and 25 mg/L chloramphenicol for strains harboring the plasmids for the target MTases. For *in vivo* libraries, non-selective medium is 2x YT or M9 media (1X M9 salts, 0.1 mM CaCl_2_, 2 mM MgSO_4_, 0.2% glucose) supplemented with 25 µg/mL chloramphenicol, 50 mg/l L-cysteine. Selective medium was M9 media with chloramphenicol, 1 mM protocatechuic acid and 1 mM methionine, without L-cysteine. Cells were made competent and transformed using TSS buffer^20^ for routine strain construction, and electroporation for library construction.

### Strains and plasmids

The selection strain SDT1065 was constructed from an in-house strain MTP3, which is derived from HMP174^10^. *CYS3* and *CYS4* were PCR amplified along with expression elements from pHM11^10^ and integrated into the chromosome of MTP3 using λ-red recombination. The full genotype of the selection strain SDT1065 is derived from *E. coli* BW25113 with FolE (T198I) YnbB (V197A) ΔtnaA ΔcysE Δcfa ompW-Δ(26 bp intergenic region)>yciE::cys3-cys4. Growth coupling with this strain was verified by growth in cysteine-free defined media with PCA and methionine supplementation, and various versions of MTase with previously known activity levels.

Plasmids pSD221 and pSD293 are used for expressing MTases, with the low-copy number pSC101 ori, Chloramphenicol acetyltransferase as the antibiotic resistance marker, and a linker for Golden Gate assembly. The linker has two BsaI digestion sites on the two ends of the linker region. The two plasmids differ in that pSD221 has J23100 while pSD293 has the weaker J23110 (Anderson Biobricks) as the expression control promoters. Both use RBSU058^21^ and phase lambda terminator as the expression control RBS and terminator.

All strains and primers used are listed in **Extended Data 3 G 4**.

### Library generation and barcoding

#### Error-prone PCR

epPCR was initially performed on wild-type *AtOMT1* codon-optimized for *E. coli* to generate the first OMT library. The second OMT library used for data collection in this work was generated by epPCR on the plasmids extracted from the selection results of the first library. GeneMorph II Random Mutagenesis Kit was used according to manufacturer’s instructions, with 1 ng of template DNA and 30 cycles. The forward primer additionally contains sequences used for the assembly of the variants onto the cloning vector in a later step. Several reactions were run in parallel to increase the amount of products. The PCR products were gel-purified to remove residual primers.

#### Barcoding with one-cycle PCR

A one-cycle PCR step was performed on the epPCR products to attach random, 25-bp barcodes to the DNA variants, using a single primer which carries the barcodes. All purified epPCR products are used in the reaction (separated into several reactions as needed). 2X Q5 hot-start master mix is used. The reaction is done with 150s of 98 °C for denaturation, 150 seconds at the annealing temperature (68 °C in this case), and 8 min of 72 °C for extension. The prolonged time for each step is to maximize the extent of the reaction for all molecules.

The one-cycle PCR product is purified through gel purification, which also serves the purpose of removing the leftover single strand in the process. The barcoded library is ligated to pSD221 via Golden Gate assembly in several parallel reactions. Afterwards, the reactions were pooled together and purified with AMPure XP beads (Beckman Coulter) according to manufacturer’s PCR purification protocol with 0.9 bead-to-reaction volume. The entire library is eluted to ∼25 µL of water.

### Construction and amplification of *in vivo* library

#### Transformation of the plasmid library

Electrocompetent cells for the selection strain were prepared according to commonly used protocols. Several electroporations were done in parallel to transform the full amount of the plasmid library. Immediately after electroporation, the cell suspensions were diluted in 2x YT media (∼1:10 volume ratio) without antibiotics and incubated for 1 hour at 37 °C for recovery.

After recovery, glycerol stocks were made of the transformant library and stored at - 80°C. A small volume of the library is serially diluted and plated onto Chloramphenicol plates to estimate the library size per volume of the stocked library.

#### *In vivo* library amplification

Given current technologies, the library size characterized by this invention is one to tens of millions per run, limited by long-read sequencing capacity. If all variants are to be profiled, aliquots of the transformant library within the expected profiling limit should be used.

An amount of the transformant library aliquots based on the consideration above is grown in 2x YT media with chloramphenicol and cysteine overnight for amplification so that aliquots of the same library can be used in subsequent steps. The amount of media should allow sufficient amplification of the library such that sufficient aliquots can be made and aliquots can remain representative of the library distribution. The amplified library is aliquoted and stocked as glycerol stocks at -80°C.

### Long-read sequencing

#### Plasmid extraction and digestion

One aliquot of the *in vivo* library was grown in a large volume of rich, non-selective media (2x YT) with chloramphenicol and cysteine at 37 °C with shaking. The plasmids were extracted and purified using maxiprep (ZymoPURE™ II Plasmid Maxiprep Kit) or miniprep (Macherey-Nagel NucleoSpin® Plasmid) according to manufacturer’s instructions. Due to limited data output from single Nanopore runs, three runs were performed on the unselected library to obtain sufficient reads.

In addition, plasmids from one post-selection culture were also processed and sequenced in one Nanopore run. This was done by using a 12 OD aliquot to inoculate 100 mL of selective medium (an initial OD of ∼0.12), which was allowed to grow until OD reached 1.28. At this point, 2x YT with cysteine were added to allow non-selective amplification before plasmid extraction.

The extracted plasmid was digested with SalI and SpeI (both from NEB), two unique restriction sites on the plasmid sequence. SalI is 71 bp upstream of the expression promoter, while SpeI is 46 bp downstream of the barcode region. The digestion reaction was gel purified to extract the expression cassette-barcode region.

#### Nanopore sequencing

This step was done with the R10.4.1 flow cell on a GridION instrument, with sample preparation done using Ligation Sequencing Kit V14 according to the protocol “Ligation sequencing amplicons V14 (SQK-LSK114)” from Oxford Nanopore. Additional materials and instruments were used according to the protocol: NEBNext® Companion Module for Oxford Nanopore Technologies® Ligation Sequencing, Qubit fluorometer and the associated consumables, AMPure XP beads and magnetic rack for 1.5 ml Eppendorf tubes, 1.5 ml Eppendorf DNA LoBind tubes.

After sample preparation, the sample was loaded onto a GridION and run for 39 hours with 9.48 M reads generated (longer run time can be done for higher read output). 100% of the reads were called, and 15.76 Gb bases were called (based on min Q score of 7, though the overall Q score is generally above 12 throughout the run).

20.8 million reads in total were obtained from Nanopore sequencing, including 19.3 million in non-selected library, and 1.5 million in selected library.

### Selective growth sampling and deep sequencing

#### Selective growth sampling

An aliquot of the library was grown overnight in non-selective medium at 37 °C with shaking. The next day, the culture was washed twice with the selection medium, and inoculated into the selection medium at an initial OD of ∼0.1, at 30 °C. Another portion of the overnight culture was spun down, washed twice with water, and the cell pellet was stored at -20 °C, which served as the sample at time point 0.

Further samples were taken at 43 hr (OD ∼0.24, denoted time point 1), 50 hr (OD ∼0.35, denoted time point 2. A dilution into an initial OD of 0.05 was made at this point to maintain exponential growth.), 67 hr (OD reading ∼0.27, or cumulative OD of ∼1.9, denoted time point 3), 71 hr (OD reading ∼0.45, or cumulative OD of ∼3.2, denoted time point 4). All were done in triplicates.

#### Sample preparation for deep sequencing

Cell pellet samples from growth-based selection were thawed, and water was added to reach a final concentration of approximately 0.3 OD per 10 µL. Thermolysis was performed by heating the cell solution at 95 °C for 10 min. The resulting mixture was centrifuged, and the supernatant was kept as PCR template for barcode amplification.

For each lysed cell pellets, barcodes were amplified as follows for Illumina sequencing using Kapa HiFi Hotstart ReadyMix according to manufacturer’s instructions. The amplicons were further barcoded with custom Illumina barcodes according to the TruSeq UDI scheme. The PCR products were purified with AMPure XP beads. The purified products were quality checked with Qubit.

#### Deep sequencing

>1 µg of PCR purified PCR product was sent to GENEWIZ Germany GmbH (Azenta Life Sciences) for Illumina sequencing. The sequencing platform was Illumina NovaSeq 2x150 bp sequencing. 20-33 million Illumina reads were obtained for each sample.

### Bioinformatics processing to determine the sequence-fitness relationship

#### Mapping Sequence to Growth Rate

Using custom scripts, Illumina reads with a mean -q score below 30 were removed. The barcodes were extracted from the reads producing a CSV file with the abundances of each unique barcode across all time points. Subsets of the dataset were generated, each containing barcode abundances at time point pairs 0-1, 0-2, 0-3, and 0-4.

The DiMSum suite^11^ was used to compute the growth rates for each of the dataset subsets with the following parameters: –maxSubstitutions 100 –mixedSubstitutions T – indels all –startStage 4 –stopStage 5. The result is a mapping of each barcode to a set of growth rates.

Nanopore reads were basecalled using the Dorado v5.0.0 sup-accuracy basecaller. Reads containing multiple concatenated coding regions were split into multiple reads using a custom script. Filtering based on read quality and read length was performed using Chopper with parameters: -q 9 -l 800 –maxlength 2000.

UMIC-seq^8^ was then used to extract the barcodes from the filtered ONT reads. Separately, barcodes extracted from Illumina reads were clustered using mmseqs2 easy-linclust^22^ with parameters –min-seq-id 0.88 -c 0.88, and the most abundant barcode in each cluster was collected in a table. Using vsearch –usearch_global with parameters –query_cov 0.84 and –id 0.88, barcodes derived from ONT reads were clustered against the barcodes in the table. For clusters containing seven or more IDs, the reads were collected in a FASTA file.

Using a custom pipeline, a consensus sequence was generated for each cluster. minimap2^23^ was used to align the reads in each cluster to the reference sequence, and racon^24^ was used to generate a draft assembly. In three additional rounds, minimap2 and racon were used with the generated draft assembly replacing the reference sequence as the template. The final draft assembly was then polished with medaka (ONT) using one round of medaka_consensus, producing a consensus sequence for each cluster.

The consensus sequences, along with their corresponding barcodes, were compiled into a CSV file. The growth rate scores were then linked to the consensus sequences using the barcode as the identifier.

#### Nanopore read count requirement evaluation

The error rate of the consensus sequence for a coding region depends in part on the number of reads in the cluster used to produce it. A lower bound for the coding region error rate can be provided based on the read count of the cluster. Self-consistency was estimated by computing the coding region of a set of 700 clusters with 25 reads or more. Having 25 reads or more in a cluster implies the consensus sequence will be highly accurate. Sub-sampling the clusters, we generated clusters ranging from 7 to 15 in size. For each cluster size between 7 and 15, we created 5 subsets. The coding regions computed from the subsets were compared to those computed from the full cluster. The frequency of mismatched coding regions was then estimated for cluster sizes 7 to 15, respectively.

#### Post Processing the Dataset

The coding region and promoter region were extracted using a custom script. The consensus sequence was aligned to the reference sequence, and all bases between the first and last base of the promoter and coding regions were extracted, respectively.

Since the template for the construction of the library contained a mixture of variants that were evolved through an initial selection step, Parent2 coding sequence in this example was defined as the most common sequence in the unselected library, containing six point mutations compared with Parent1 (see full sequences in **Extended Data 8**).

We created an aggregated version of the dataset in which, for each variant 𝑣, an aggregated growth rate 𝜆_𝑣_− is produced from the set of growth rates. Using the equation below,

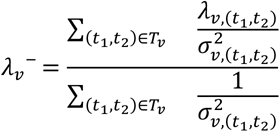

the aggregated growth rate 𝜆_𝑣_− for a variant 𝑣 was computed using only the time-step pairs 𝑇_𝑣_ for which the variant has both 𝜆_𝑣,(𝑡,𝑡)_ and 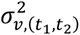 values.

#### Unique protein sequence enumeration

First, the following variants were removed from the original dataset (data_seq_to_fitness_t_0_x_no_vif.csv): 1) variants with a stop codon, 2) variants with frameshift mutations (indels) in the DNA coding region, 3) variants with any mutations (substitution or indel) in the promoter region, 4) variants with indels in the protein sequence. Then, for every unique protein sequence, growth rates from all degenerate nucleotide sequences were averaged to obtain the column growthrate_total. If multiple nucleotide sequences were present, growthrate_sigma_total was set to the standard deviation of each growthrate_total value per nucleotide sequence. If only one nucleotide sequence was present for the protein sequence, then the nucleotide growthrate_sigma_total was used.

### Learning from the data to generate sequence variants

#### Training a Second-Order Model Using MoCHI

MoCHI was used to fit a second-order model predicting variant growth rates based on additive effects of individual mutations and pairwise epistatic interactions. The model training parameters were set to –min_observed 4, –max_interaction_order 2, – learn_rate 0.02, and –transformation ReLU, with 10-fold cross-validation. A filtered version of the non-aggregated dataset, corresponding to time step pair 0-4, was used as the training set to ensure an accurate measure of variance. Variants containing indels, promoter region changes, or stop codons before the last active site were excluded, resulting in a dataset of 103,731 records. MoCHI provides a score for each single mutation and pairwise epistatic interaction inferred from the training set, as well as a score for Parent2. Using the defined parameters, a positive score indicates a beneficial effect on the growth rate, while a negative score indicates a detrimental effect.

#### Generating Variants Using the Second-Order MoCHI Model

Scores obtained for mutations and pairwise epistatic interactions were used to generate a library of high-growth-rate variants. A greedy iterative sampling algorithm was employed to identify a set of mutations that, when applied to Parent2, maximize the predicted growth rate according to the MoCHI model. To generate variants, we optimized for the sum of additive traits. A function 𝑓 is defined as the sum of the mean single mutation terms <u>𝑆</u> and the mean pairwise epistasis terms <u>𝐸</u>, as shown in equations below:

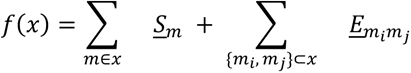

Since the second-order model was trained using ten-fold cross-validation, up to ten different scores are available for each single mutation or pairwise epistasis term.

The sampling algorithm operates in two phases. In the first phase, an empty set of mutations (the nominated set) is initialized. During each iteration, a position not yet occupied by a mutation in the nominated set is randomly sampled. All mutations with an available score (candidate mutations) are then evaluated using equation below:

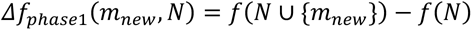

From the set of candidates with a positive 𝛥𝑓_𝑝ℎ𝑎𝑠𝑒1_ (beneficial mutations), one mutation was sampled and added to the nominated set. The probability distribution for sampling was determined by applying the softmax function to the 𝛥𝑓_𝑝ℎ𝑎𝑠𝑒1_ values of all beneficial mutations, using a temperature 𝑇_𝑚𝑎𝑥_. This process was repeated until the desired number of mutations was reached.

In the second phase, mutations in the nominated set are iteratively removed and replaced with better ones, if available. During each iteration, a position not yet occupied by a mutation in the nominated set is randomly sampled. All possible combinations of adding a candidate mutation and removing an existing one from the nominated set are evaluated using equation below:

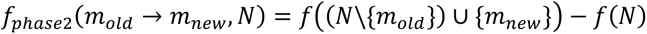

For each candidate 𝑚_𝑛𝑒w_, the mutation swap 𝑚_𝑜𝑙𝑑_ → 𝑚_𝑛𝑒w_ with the highest 𝛥𝑓_𝑝ℎ𝑎𝑠𝑒2_was added to the set of beneficial swaps, provided 𝛥𝑓_𝑝ℎ𝑎𝑠𝑒2_ was greater than 0. From this set, one swap was sampled, adding 𝑚_𝑛𝑒w_ and removing 𝑚_𝑜𝑙𝑑_ from the nominated set. The probability distribution was determined by applying the softmax function to the 𝛥𝑓_𝑝ℎ𝑎𝑠𝑒2_ values of all beneficial swaps. Sampling was performed using a temperature 𝑇_𝑖_, which followed a linear schedule from 𝑇_𝑚𝑎𝑥_ to 𝑇_𝑚𝑖𝑛_. The iteration process stopped when for all positions no more beneficial swaps were available or the maximum number of iterations, 𝐼_𝑚𝑎𝑥_, was reached.

A subset of the generated variants was selected for synthesis. Variants with a negative pairwise epistatic interaction term below a threshold - set at half the negative score of Parent2 - were excluded. Additionally, variants were removed if less than one-third of their potential epistatic effects were accounted for by the MoCHI model. From the filtered pool, those offering the best balance of diversity and growth rate were selected for synthesis.

We generated 200 sequences each with 7 and 15 mutations using 𝐼_𝑚𝑎𝑥_ = 1000, and 50 sequences each with 25 and 35 mutations using 𝐼_𝑚𝑎𝑥_ = 1500. The temperature parameters, 𝑇_𝑚𝑎𝑥_ and 𝑇_𝑚𝑖𝑛_, were set to 5 and 0.5, respectively. Duplicate sequences were removed, and filters were applied, reducing the total number of sequences to 115. We selected 3 sequences each with 7 and 15 mutations, and 2 sequences each with 25 and 35 mutations.

To convert amino acid sequences to DNA sequences for synthesis, we used the coding region of Parent2 as the template, and introduced the amino acid mutations using the most frequent codon for each desired amino acid in E. coli.

#### Deterministic Naive Variant Generation Using the Second-Order Model

As a comparison, we generated a library with minimal consideration of epistasis. Sequences were generated by sequentially adding mutations from the set of available mutations, in decreasing order of score, to the nominated set until the desired number of mutations was reached. The mutation set included all single mutations and pairs provided by the MoCHI model. Single mutations were assigned the score provided by MoCHI, while pair scores were calculated by combining the individual mutation scores and the epistatic term, based on the equation below:

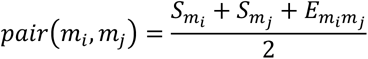

Pairwise epistasis with mutations already in the nominated set was ignored. When a pair was selected, both mutations were added to the set. In the final iteration, only single mutations were considered.

### Experimental validation of the data and the generated sequences

#### Strain construction

34 AtOMT1 variants evolved during growth-based selection with different estimated fitness levels were selected for validation experiments (see **Extended Data 5** – all variant sequences). All variants had consensus sequences based on ONT reads. By using forward primer SD_PR757 (from start codon) and a reverse primer annealing to the unique barcode for the variant of interest, variants were individually PCR amplified from the miniprepped plasmids of non-selected or selected libraries. Reverse primers used for isolating variants from the mixed template were SK_PR47-SK_PR72, SK_PR115, and SK_PR128-SK_PR148. Amplified variant coding regions were gel purified and assembled into plasmid pSD293 using Golden Gate. The promoter and coding region of each resulting plasmid were verified by Sanger sequencing. In all cases, the sequence of the cloned variant matched the deep sequencing-based consensus sequence with 100% identity.

For growth assays, strains were constructed by transforming the pSD293-derived plasmids into the selection strain SDT1065. For bioconversion assays, the plasmids were transformed directly into E. coli BW25113 instead to avoid any contribution of the unique variants to fitness of when generating biomass for bioconversion. All 34 variants were successfully cloned into SDT1065, and 32 of the 34 variants were successfully cloned into BW25113.

The barcodes of evolved variants were preserved during the cloning process. A random, unique barcode was attached to Parent1 using reverse primer SD_PR758. As with the evolved variants, the barcoded Parent1 was cloned into pSD293, resulting in both BW25113 and SDT1065-derived strains. An additional BW25113-based Parent1 strain was constructed with the barcode of the high-fitness evolved variant 17 attached to the end of the Parent1 coding sequence by using primers SD_PR757 and SK_PR172. This was done to control for the effect of the barcode on expression and mRNA stability.

Variants generated using MoCHI were ordered as gene fragments from Twist Bioscience. 22 out of out of the 26 generated variants (variants with prefixes o1_ and o2_ in **Extended data 5 G 6**) were successfully cloned into pSD293 in BW25113 for the bioconversion assay and verified by Sanger sequencing. o2_t7 and o2_t25 had a G2D mutation and a D2G mutation, respectively. These two variants are included in the presented bioconversion assay results (**Fig. 2F**) with an asterisk (*) to indicate the deviation from the original design.

#### Growth assay

SDT1065 derivatives expressing variants of AtOMT1 under the control of J23110 in pSD293-derived plasmids were grown overnight in 2xYT media supplemented with cysteine supplemented with 25 µg/mL chloramphenicol. These precultures were washed with equal volume of 0.85% NaCl. In triplicates for each strain, 5 µL of the washed precultures were inoculated into 245 µL of selective medium. The cultures were grown in a Growth Profiler (Enzyscreen BV, The Netherlands) and growth was monitored over time. Growth rate was estimated with the built-in software of Growth Profiler. Growth rates were calculated individually for each replicate. A mean growth rate was calculated for each strain.

#### Bioconversion assay

*E. coli* BW25113 strains harboring variants of AtOMT1 cloned into pSD293 were grown overnight in 2xYT media supplemented with 25 µg/mL chloramphenicol in 96-deepwell plates placed in a MaxQ 8000 incubator set to 30 °C and 300 rpm. On the next day, each overnight culture was used to inoculate six new 500 µL cultures in new 96-deepwell plates with 2xYT supplemented with chloramphenicol to an initial OD of 0.05. The new plates were placed back in the MaxQ 8000 incubator at 30 °C, 300 rpm.

After 4 hours, the OD_600_ of the cultures was measured to be between 0.54 and 1.51. The plates were put on ice, and cell mass was normalized to 1 mL of cells with an OD_600_ of 0.54 in triplicates for each strain by combining culture from two wells for each triplicate. Because each preculture was used to inoculate six 500 µL cultures, the triplicates could be kept independent by transferring culture from two new wells for each triplicate. New plates with combined, normalized samples then centrifuged at 1500 × g for 15 minutes to pellet the cells. The supernatant was removed, and the cell pellets were stored at -70 °C overnight.

Next day, the cell pellets were equilibrated to room temperature, and 70 µL of room-temperature B-PER Complete Bacterial Protein Extraction Reagent (B-PER) (Thermo Scientific) was then added to each pellet. The resuspended pellets were incubated at room temperature with gentle rocking (40 rpm) for 30 minutes.

After lysis, 50 µL of B-PER lysate was mixed with 50 µL of bioconversion buffer consisting of an aqueous solution of 1 mM PCA, 2 mM L-methionine and 2 mM ATP disodium salt (A2383, Sigma-Aldrich). The 100 µL bioconversion reactions were incubated in sealed microtiter plates for 120 minutes at 30 °C and 246 rpm in a Labnet 311DS Environmental Shaking Incubator. Reactions were stopped by heating the microtiter plates in a 95 °C water bath for 5 minutes.

The microtiter plates were centrifuged at 1000 × g for 15 minutes to pellet cell debris. The supernatant was filtered into HPLC sample plates through AcroPrep™ Advance Plates, Short Tip (Cytiva, 97052-096) and diluted 2x with Blank Buffer to ensure sufficient volume for HPLC. Blank Buffer components were 0.5 mM PCA, 1 mM L-methionine, 1 mM ATP disodium salt, and 50% B-PER.

Samples were analyzed on an Ultimate 3000 HPLC with a Zorbax C18 column coupled to a DAD-3000 module. The sample injection volume was 10 µL, and the eluents were 0.05% acetic acid and 100% acetonitrile. The amount of acetonitrile in the flow gradient was initially 5%, increasing linearly to 30% at 8.75 minutes, 40% at 11.00 minutes, and 60% at 12.5 minutes. The amount of acetonitrile stayed at 60% until 13.50 minutes, where it decreased linearly to 5% at 14.00 minutes. The flow rate was constant at 1.0 ml/min. The length of the method was 15 minutes. Isovanillic acid was observed as a peak in absorbance at 260 nm at a retention time of 5.823 minutes.

For quantification, an isovanillic acid standard was analyzed along with the samples. Isovanillic acid was dissolved in Blank Buffer to a concentration of 487.55 mg/L, and a dilution series was made using Blank Buffer to generate concentrations of isovanillic acid between 6.424 µg/L and 487.55 mg/L. The linear dynamic range was found to be between 31.2 µg/L and 97.51 mg/L. The 2X dilution of samples with blank buffer after filtration was accounted for when presenting the concentration of isovanillic acid in the samples.

For the PCA bioconversion assay, variant activities were measured in triplicate. Because replicate 1 consistently yielded lower isovanillic acid concentrations across most variants, we inspected potential systematic effects. Biomass for each replicate was grown in separate deep-well plates, and all replicate 1 cultures originated from the same two plates. We therefore attribute the discrepancy to plate-specific growth conditions. Consequently, replicate 2 and 3 were used for the final analysis and for scatterplots correlating fitness with isovanillic acid production.

For Var12, bioconversion replicates 1 and 2 showed no detectable isovanillic acid, while replicate 3 yielded 0.268 mg/L. Because variant 12 carries a premature stop codon and had shown no product formation in previous independent bioconversion assays, replicate 3 was considered contaminated and excluded from further analysis. Only replicate 2 (0 mg/L) was used for the scatterplot in **Fig. 2B** (see **Extended Data 9** for all measurements).

---

**Supplementary Figure 1:**
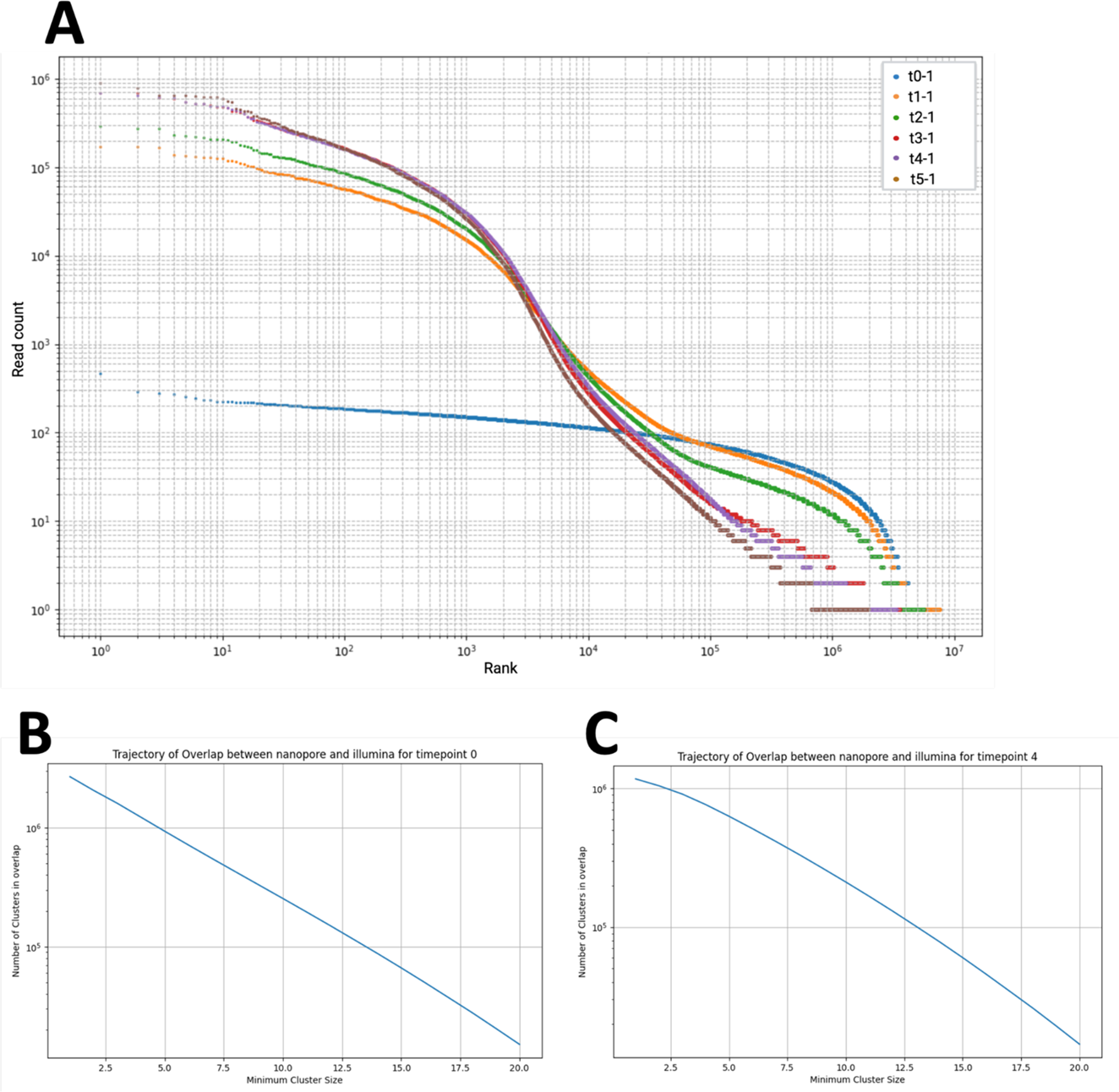
Barcode count dynamics. **a**, Individual barcode read count distribution for replicate 1 over 6 sampling time points. Over the course of selection, the distribution becomes less even. **b**, Barcode overlap between Nanopore and Illumina data at time point 0. **c**, Barcode overlap between Nanopore and Illumina data at time point 4, being slightly lower than that of time point 0.

**Supplementary Figure 2:**
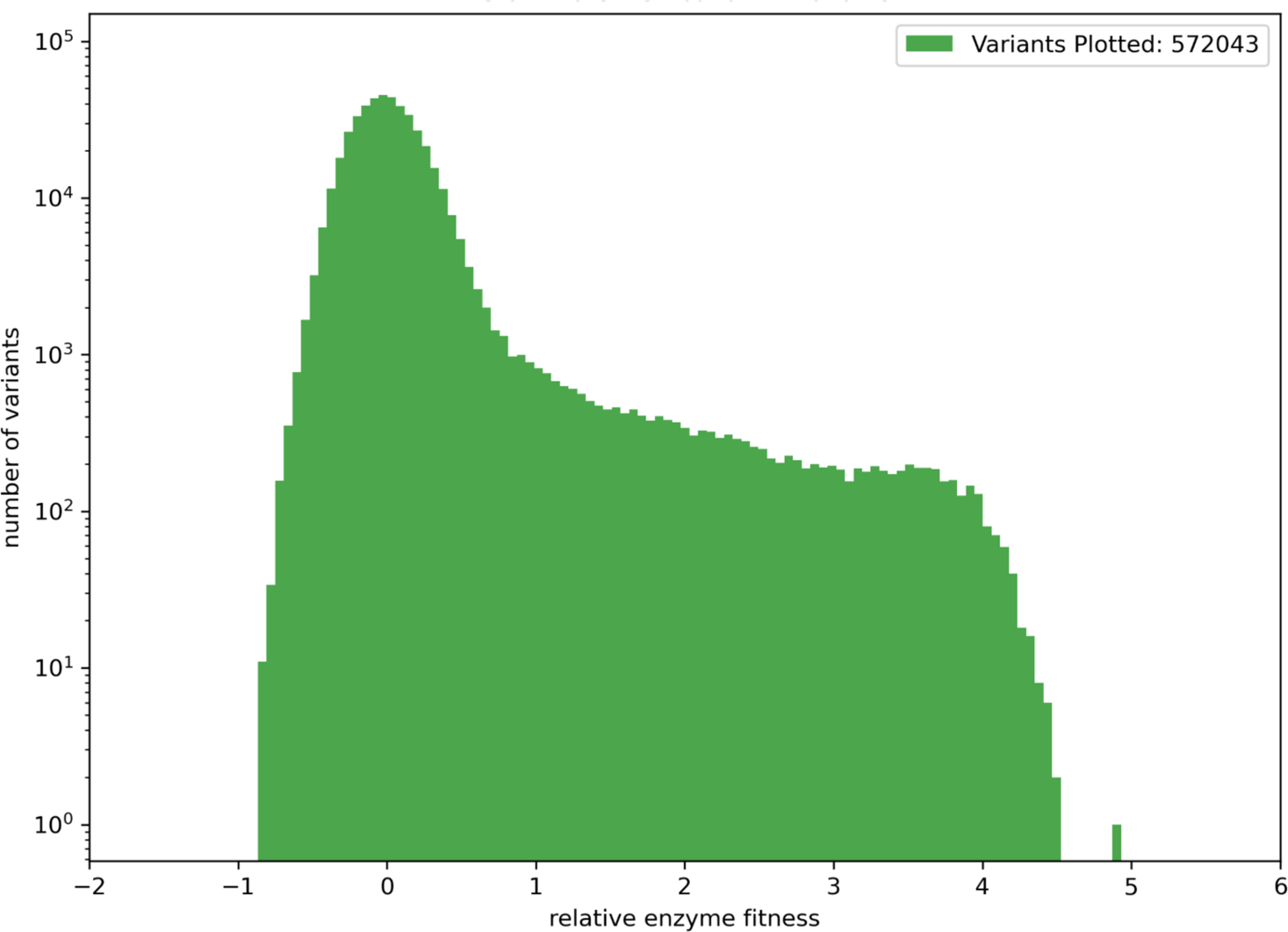
The scale and fitness range of the extracted MillionFull sequence-fitness data for AtOMT1. Variant fitness levels were obtained by normalizing variant growth rates with the average growth rate of variants with Parent2 sequence.

**Supplementary Figure 3:**
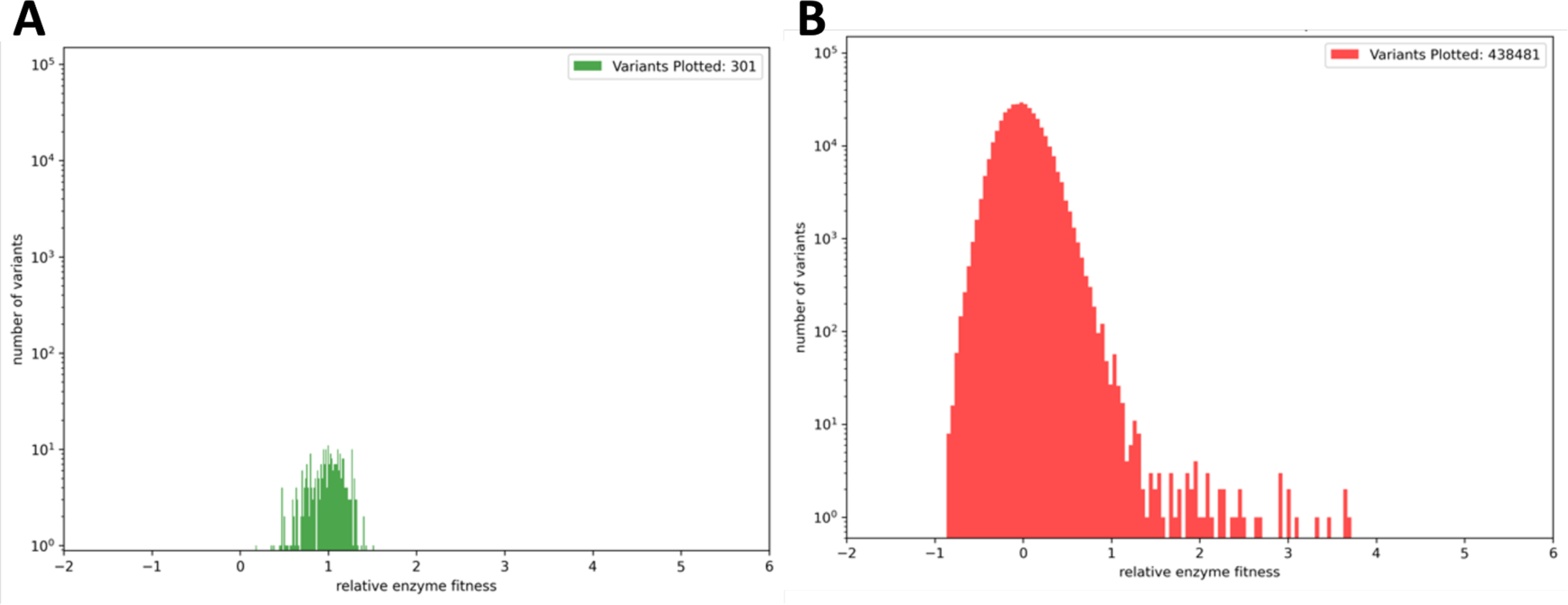
Sanity data check. **a,** Fitness levels of variants with Parent2 sequence. **b,** Distribution of fitness levels for variants with premature stop codons.

**Supplementary Figure 4:**
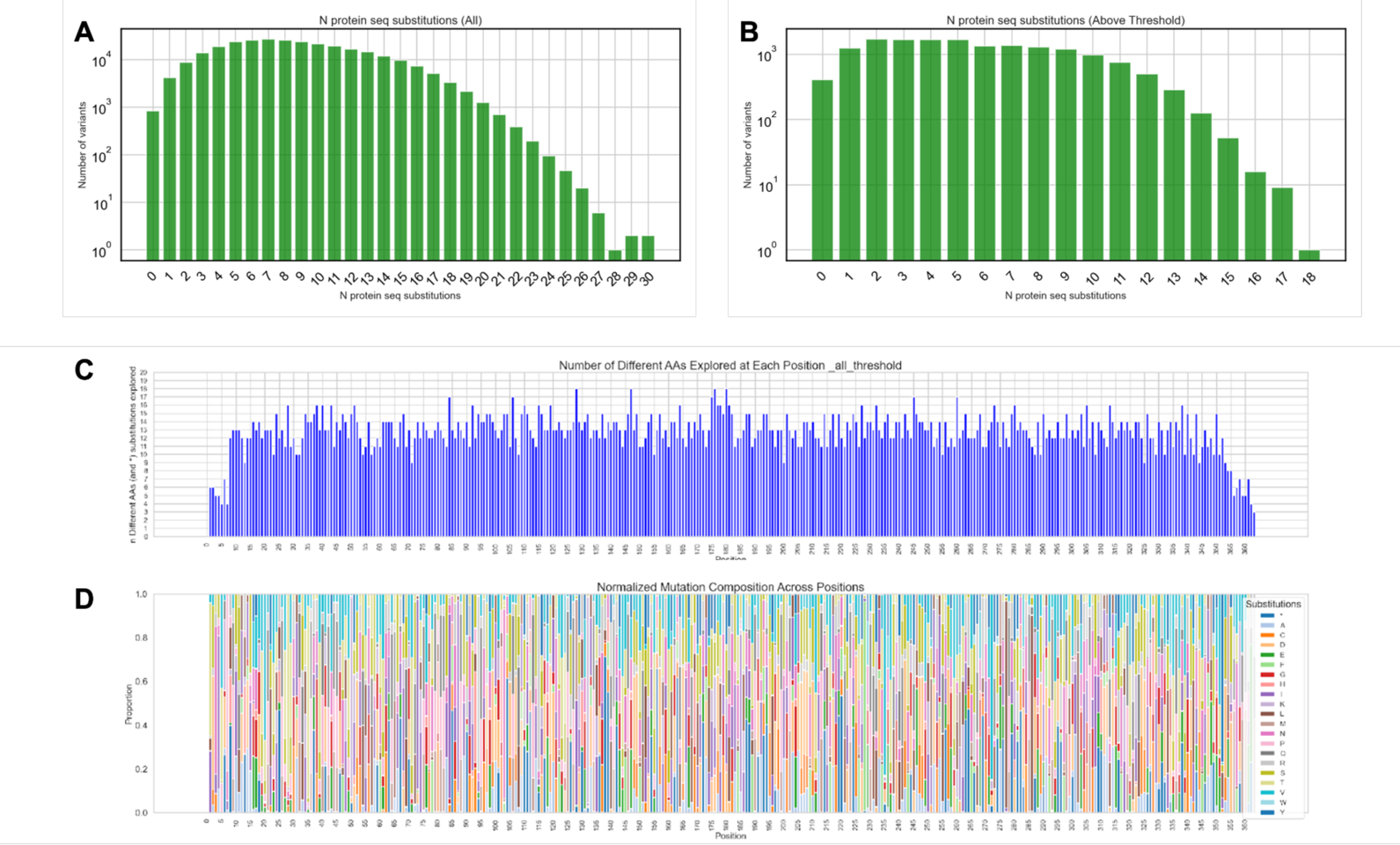
Mutation distribution across variants in the collected MillionFull dataset. **A** Distribution of number of amino acid substitutions for variants without premature stop codons or indels. **B** Distribution of the number of amino acid substitutions for variants with growthrate higher than Parent2 by 0.05. **C** Number of amino acids substitutions present at each position in the collected MillionFull dataset. **D** Composition of mutations at each position in the collected MillionFull dataset.

**Supplementary Figure 5:**
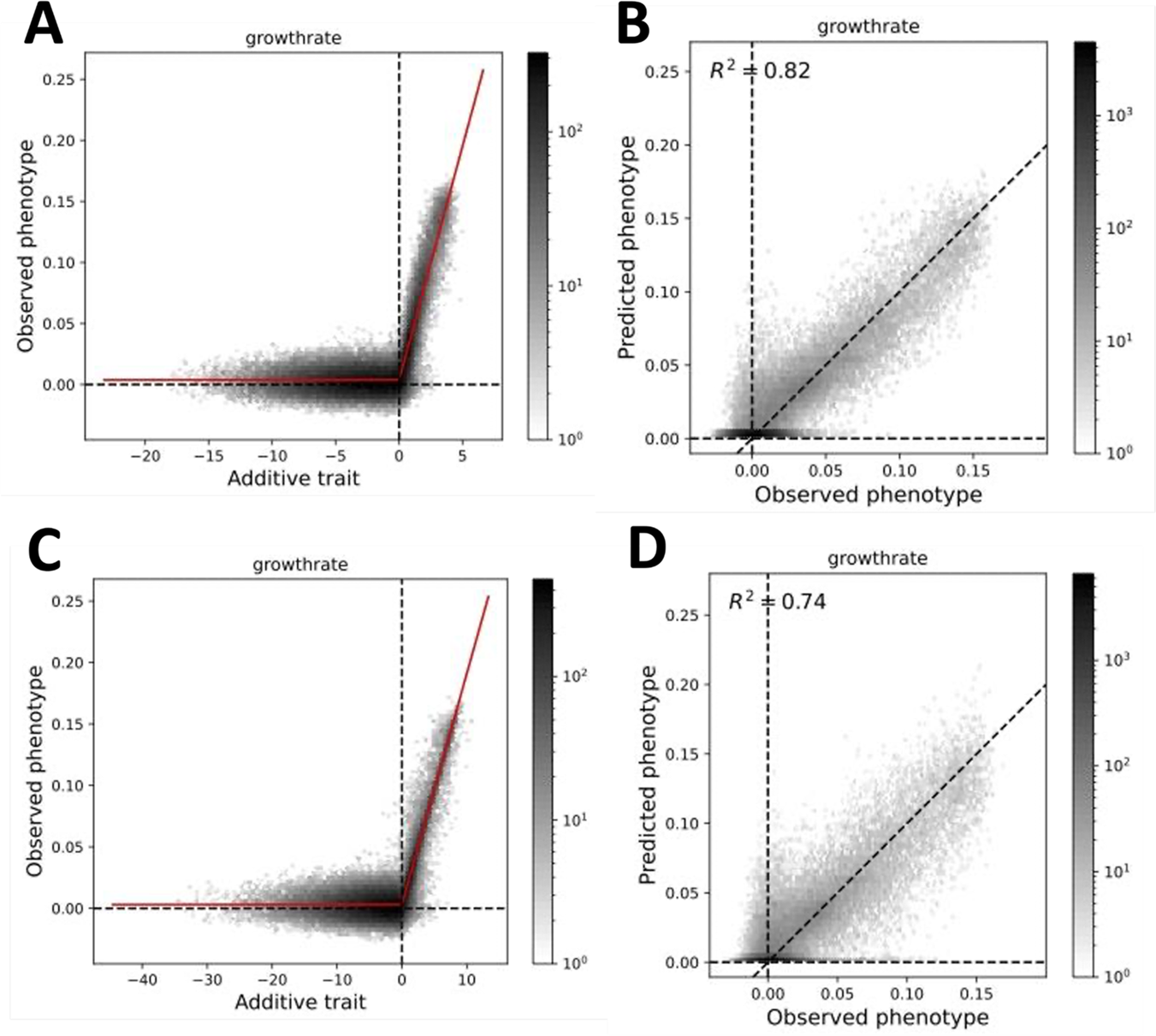
MoCHI model fit quality. A, **B** First order model.**C, D** Second order model. See Ref. 13 for detailed interpretation.

**Supplementary Figure 6:**
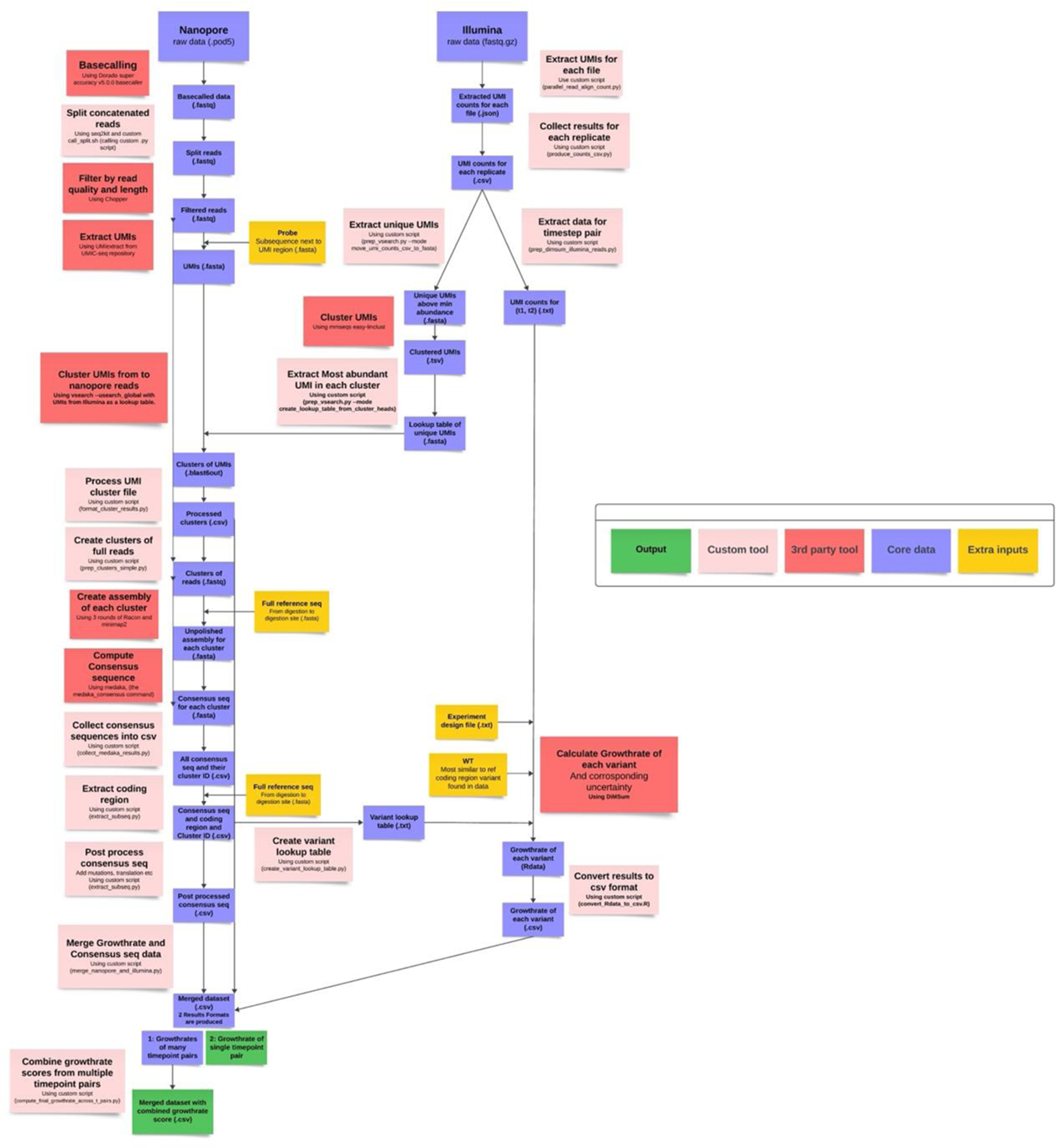
The MillionFull bioinformatics pipeline.

