## Extended data-MillionFull for "MillionFull enables low-cost, massive, full-length enzyme sequence–fitness data collection for machine learning–guided enzyme engineering": Extended data 1 - t_0_4 dimsum report.html


### DiMSum Report


#### Settings

**DiMSum version:** 1.3.1

**Project name:** DiMSum\_Project

**Run Started:** 2024-10-27 23:02:50

**Run Completed:** 2024-10-28 01:09:38

**Command-line arguments:**

```
## runDemo                 FALSE
## fastqFileExtension      .fastq
## gzipped                 TRUE
## stranded                TRUE
## paired                  TRUE
## barcodeErrorRate        0.25
## experimentDesignPath    data/Illumina/ASMT/intermediate/DiMSum/ExperimentDesign_ASMT_timepoints_0_4.txt
## experimentDesignPairDuplicates
##                         FALSE
## countPath               data/Illumina/ASMT/intermediate/DiMSum/variantCounts_ASMT_timepoints_0_4_merged_identical_variants.txt
## cutadaptMinLength       50
## cutadaptErrorRate       0.2
## cutadaptOverlap         3
## vsearchMinQual          30
## vsearchMaxQual          41
## vsearchMaxee            0.5
## vsearchMinovlen         10
## outputPath              data/Illumina/ASMT/intermediate/DiMSum/AnalysisResults_no_weighing_no_vif_aggregated_variants_timepoint_0_4
## projectName             DiMSum_Project
## wildtypeSequence        GAGTAGCTACTAGAAATCGTAGAGT
## reverseComplement       FALSE
## sequenceType            auto
## mutagenesisType         random
## transLibrary            FALSE
## transLibraryReverseComplement
##                         FALSE
## bayesianDoubleFitness   FALSE
## bayesianDoubleFitnessLamD
##                         0.025
## fitnessMinInputCountAll
##                         0
## fitnessMinInputCountAny
##                         0
## fitnessMinOutputCountAll
##                         0
## fitnessMinOutputCountAny
##                         0
## fitnessHighConfidenceCount
##                         10
## fitnessDoubleHighConfidenceCount
##                         50
## fitnessNormalise        TRUE
## fitnessErrorModel       TRUE
## indels                  all
## maxSubstitutions        100
## mixedSubstitutions      TRUE
## retainIntermediateFiles
##                         TRUE
## splitChunkSize          3758096384
## retainedReplicates      all
## startStage              4
## stopStage               5
## numCores                1
```

#### Pipeline stages

The DiMSum pipeline consists of five stages grouped into two modules
which can be run independently:

- **WRAP** (Stages 1-3) processes raw FastQ files
  generating a table of variant counts
- **STEAM** (Stages 4-5) analyses variant counts
  generating variant fitness and error estimates

Below you will find summary plots with results of each stage
corresponding to the module(s) that were run.

#### 4. **PROCESS** variants (STEAM)

DiMSum Stage 4 (PROCESS) processes sequences and filters them in
order to retain user-specified nucleotide or amino acid substitution
variants of interest. The result is a table of variant counts for all
samples. Read count diagnostic plots can then be used to rapidly check
for the presence of problematic variants (likely the result of
sequencing errors) and take steps to remove them (see Sections 4.1 and
4.2 below).

The plot below shows the **percentage of reads**
retained or discarded in each sample according to the following
criteria:

- **‘0 hamming dist.’** (retained: wild-type
  sequence)
- **‘1 hamming dist.’** (retained: 1 nucleotide
  substitutions from wild-type sequence)
- **‘2 hamming dist.’** (retained: 2 nucleotide
  substitutions from wild-type sequence)
- **‘3+ hamming dist.’** (retained: >3 nucleotide
  substitutions from wild-type sequence)
- **‘indel’** (retained: insertion or deletion
  variant)
- **‘mixed’** (discarded: nonsynonymous variants have
  synonymous substitutions in other codons, see ‘mixedSubstitutions’
  option)
- **‘too many’** (discarded: too many nucleotide or amino
  acid substitutions, see ‘maxSubstitutions’ option)
- **‘not permitted’** (discarded: nucleotide substitution
  not permitted, see ‘wildtypeSequence’ option)
- **‘internal constant region’** (discarded: nucleotide
  substitution within internal constant sequence, see ‘wildtypeSequence’
  option)
- **‘indel discarded’** (discarded: insertion or deletion
  variant)
- **‘invalid barcode’** (discarded: reads represent
  barcode sequences, but are not found in the user-supplied barcode
  identity file, see ‘barcodeIdentityPath’ option)

**Note**: The plots below show read counts
**before** application of user-specified count
thresholds.

See DiMSum
documentation for more details.

Nucleotide variant statistics (counts). The plot below is similar to
the one above instead the **total number of reads** (rather
than the percentage) in each sample is shown.


##### 4.1 Input count distributions

The **diagnostic plot** below shows marginal variant
count distributions separately for all Input samples and stratified by
the number of nucleotide substitutions (Hamming distance to the
wild-type sequence). Distributions corresponding to Hamming distances
greater than 12 are not shown. Wild-type sequence counts are indicated
by the black vertical dashed line.

Expected counts from ‘fake’ variants (due to base-call errors at a
rate corresponding to the ‘vsearchMinQual’ option) are indicated by
coloured dashed lines. Bimodal distributions (or unimodal distributions
not surpassing the indicated thresholds) indicate variants originating
from sequencing errors likely due to a library ‘bottleneck’. A minimum
input count threshold should be chosen to remove such variants (see
‘fitnessMinInputCountAll’ option applied in Stage 5 and DiMSum documentation for
more details.).

**Note**: The plot below shows variant counts
**before** application of user-specified count
thresholds.

##### 4.2 Sample count correlations

The **diagnostic plot** below is a scatterplot matrix
depicting correlations between variant counts from all Input and Output
samples. Matrix cells in the upper triangle show Pearson correlation
coefficients. Matrix cells in the lower triangle show scatterplot
equivalents (hexagonal heatmaps of 2d bin counts). Matrix diagonal cells
indicate count densities.

Distinct variant populations or ‘flaps’ i.e. subsets of variants that
appear at high counts in one replicate but at low counts in another (and
not due to selection) indicate replicate or DNA extraction
‘bottlenecks’. Minimum input and/or output count thresholds should be
chosen to remove such variants (see ‘fitnessMinInputCountAll’,
‘fitnessMinInputCountAny’, ‘fitnessMinOutputCountAll’ and
‘fitnessMinOutputCountAny’ options applied in Stage 5 and DiMSum documentation for
more details).

**Note**: The plot below shows variant counts
**before** application of user-specified count
thresholds.

#### 5. **ANALYSE** counts (STEAM)

##### 5.1 Input threshold for error model

The plot below shows the minimum Input count threshold (black
vertical dashed line) above which variants are expected to span the full
fitness range (y-axis spread). Variants surpassing this threshold are
used to fit the error model. Subplots indicate data corresponding to
independent biological replicates.

A strong dependency between Input variant counts and fitness
(negative correlation) can indicate a harsh selection i.e. a high level
of variants that ‘drop-out’ or are undetected in the Output.

##### 5.2 Replicate fitness distributions

The plot below shows replicate fitness distributions (without
normalisation between replicates). The fitness of the wild-type sequence
is indicated by the vertical dashed line (Fitness=0).

Linear differences between replicate fitness distributions can be
corrected by (scale and shift) normalisation before error model fitting
(see ‘fitnessNormalise’ option).

**Note**: Only variants retained during error model
fitting are depicted i.e. those variants detected in
**all** input and output replicates and with read counts
above the corresponding minimim threshold in **all** input
replicates (see Section 5.1 above).

The plot below shows replicate fitness distributions after
normalisation. Remaining differences in the shapes of the fitness
distributions after normalisation indicate systematic errors between
replicates. Affected replicates should be excluded from error model
fitting (see ‘retainedReplicates’ option).

##### 5.3 Replicate fitness correlations

The **diagnostic plot** below is a scatterplot matrix
depicting correlations between fitness estimates from all replicates
after normalisation (see Section 5.2). Matrix cells in the upper
triangle show Pearson correlation coefficients. Matrix cells in the
lower triangle show scatterplot equivalents (hexagonal heatmaps of 2d
bin fitness). Matrix diagonal cells indicate fitness densities. Only
variants retained during error model fitting are depicted (see Section
5.1 above).

Poor correlations indicate systematic errors between replicates.
Affected replicates should be excluded from error model fitting (see
‘retainedReplicates’ option).

**Note**: Only variants detected in **all**
input and output replicates are depicted.

##### 5.4 Fitness error model

The upper panels in the plot below indicate multiplicative (upper
left panel) and additive (upper right panel) error terms estimated by
the DiMSum error model. Dots give mean and error bars indicate the
standard deviation of parameters over 100 bootstraps.

The lower left panel in the plot below shows variance of fitness
scores between replicates as a function of sequencing count-based
(Poisson) variance expectation (average across replicates). The black
dashed line indicates perfect correspondence (i.e. Y=X).

The full DiMSum error model (red line) describes deviations from the
null expectation (black dashed line) in the observed variance of fitness
scores. A good fit of the DiMSum error model (red line) to the mean
empiricial variance per bin (blue points) indicates that the estimated
parameters (upper panels) accurately describe the variation in fitness
between replicates.

**Note**: Only variants retained during error model
fitting are depicted i.e. those variants detected in
**all** input and output replicates and with read counts
above the corresponding minimim threshold in **all** input
replicates (see Section 5.1 above).

The lower right panel in the plot below compares the full DiMSum
error model (red line) to variance contributions using either Input
multiplicative error terms (cyan line), Output multiplicative error
terms (magenta line) or additive error terms (green line) only.
Additionally, dashed cyan and magenta lines indicate purely sequencing
count-based variance expectation corresponding to Input and Output
samples respectively (i.e. corresponding multiplicative error terms set
to 1).

The quantile-quantile (Q-Q) plots below assesses the performance of
the fitness error model using leave-one-out cross-validation.

Error models are trained on all but one replicate and z-scores of the
differences in fitness scores between the training set and the remaining
test replicate are calculated (i.e. fitness score differences normalised
by estimated error in training set and test replicate). Because fitness
scores from replicate experiments should only differ by random chance,
if the error models estimate the error magnitude correctly, z-scores
should be normally distributed (i.e. Y=X).

Differences between replicate z-score distributions can indicate
systematic errors. Consider excluding (outlier) replicates from error
model fitting (see ‘retainedReplicates’ option).

**Note**: Only variants retained during error model
fitting are depicted i.e. those variants detected in
**all** input and output replicates and with read counts
above the corresponding minimim threshold in **all** input
replicates (see Section 5.1 above).
